# Streaming histogram sketching for rapid microbiome analytics

**DOI:** 10.1101/408070

**Authors:** Will P. M. Rowe, Anna Paola Carrieri, Cristina Alcon-Giner, Shabhonam Caim, Alex Shaw, Kathleen Sim, J Simon Kroll, Lindsay J. Hall, Edward O. Pyzer-Knapp, Martyn D. Winn

## Abstract

**Motivation:** The growth in publically available microbiome data in recent years has yielded an invaluable resource for genomic research; allowing for the design of new studies, augmentation of novel datasets and reanalysis of published works. This vast amount of microbiome data, as well as the widespread proliferation of microbiome research and the looming era of clinical metagenomics, means there is an urgent need to develop analytics that can process huge amounts of data in a short amount of time.

To address this need, we propose a new method for the compact representation of microbiome sequencing data using similarity-preserving sketches of streaming k-mer spectra. These sketches allow for dissimilarity estimation, rapid microbiome catalogue searching, and classification of microbiome samples in near real-time.

**Results:** We apply streaming histogram sketching to microbiome samples as a form of dimensionality reduction, creating a compressed ‘histosketch’ that can be used to efficiently represent microbiome k-mer spectra. Using public microbiome datasets, we show that histosketches can be clustered by sample type using pairwise Jaccard similarity estimation, consequently allowing for rapid microbiome similarity searches via a locality sensitive hashing indexing scheme. Furthermore, we show that histosketches can be used to train machine learning classifiers to accurately label microbiome samples. Specifically, using a collection of 108 novel microbiome samples from a cohort of premature neonates, we trained and tested a Random Forest Classifier that could accurately predict whether the neonate had received antibiotic treatment (95% accuracy, precision 97%) and could subsequently be used to classify microbiome data streams in less than 12 seconds.

We provide our implementation, Histosketching Using Little K-mers (HULK), which can histosketch a typical 2GB microbiome in 50 seconds on a standard laptop using 4 cores, with the sketch occupying 3000 bytes of disk space.

**Availability:** Our implementation (HULK) is written in Go and is available at: https://github.com/will-rowe/hulk (MIT License)

## 1. Introduction

The global corpus of microbiome sequence data is being augmented daily with vast volumes of data, particularly as a result of large-scale sequencing initiatives such as the Human Microbiome Project (HMP) (Human Microbiome Project Consortium, 2012), the Earth Microbiome Project (Thompson *et al.* 2017) and Global Ocean Survey (Rusch *et al*., 2007). Data outputs will continue to increase, particularly as metagenomics within the clinical field is more widely being accepted and adopted (Mulcahy-O’Grady and Workentine, 2016), and the continued decline in sequencing costs (Forbes *et al*., 2018).

We are now at the point where our ability to analyse microbiome data quickly and effectively is the main bottleneck in our workflows, particularly when it comes to real-time sequencing platforms (Greninger *et al*., 2015; Forbes *et al*., 2018). In addition, we also need to ensure that existing microbiome data remains accessible and usable (including for end-users e.g clinicians), so that it can be readily incorporated into our new analyses, and generate testable hypotheses for validation/confirmation in experimental systems. It is becoming clear that current microbiome analytics are not suitable in this age of ‘big data’, particularly in terms of data retrieval and sample classification (Kakkanatt *et al*., 2018).

Current microbiome analytics can be largely split into referenced-based or *de novo* approaches (Morgan and Huttenhower, 2012). Whereas reference based analyses (such as taxonomic classification) can often result in large amounts of sequencing data being excluded and high computational requirements, *de novo* approaches circumvent these issues. For example, the pairwise comparison of k-mer spectra is a *de novo* analysis method that has been routinely used in recent years for clustering microbiomes using dissimilarity measures (Dubinkina *et al*., 2016; Benoit *et al*., 2016). These measures are used to identify microbiome composition changes in studies that involve longitudinal sampling or multiple isolation sites (Anvar *et al*., 2014). However, k-mer spectra can still take considerable time to compute, are relatively large in file size and new sample comparisons require additional computation. As well as this, Machine Learning (ML) frameworks will struggle to use these *de novo* outputs as feature vectors due to their scale. This is a potential barrier to the use of these methods in microbiome analytics as ML can be used to help solve many of the data problems encountered in genomics and holds great potential for microbiome analytics (Libbrecht and Noble, 2015).

The application of other dimensionality reduction techniques to genomic data address some of these issues. These techniques have ranged from distributed string mining of informative k-mers (Seth *et al*., 2014), to the recent use of Locality Sensitive Hashing (LSH) (Ondov *et al*., 2016; Luo *et al*., 2018; Brown and Irber, 2016; Rowe and Winn, 2018). MinHash is one form of LSH that has greatly improved genomic analysis speeds for operations such as sample clustering, database searching and phylogenetic estimation; it works through reducing sequence data to small, representative sketches using a set of minimum k-mer hash values (Ondov *et al*., 2016). However, although MinHash-based tools like MASH and sourmash can be used to great effect for certain microbiome analytics (e.g. what genomes are in my microbiome), there remain limitations to standard MinHash techniques; such as the loss of k-mer frequency information and the impact of relative set size on Jaccard similarity estimates (Koslicki and Zabeti, 2017; Wu *et al*., 2017). With this in mind, we assert that current *de novo* microbiome analysis methods do not enable the rapid similarity, indexing and classification operations that are required in this era of big data in microbiome research. This is particularly pertinent within a clinical metagenomics setting, as accurate and ‘useful’ data is required for downstream analysis, and clinical decision making e.g. antibiotic treatment choices (Kakkanatt *et al*., 2018).

We present a method to reduce microbiome sequence data streams to an updateable ‘histosketch’ of the underlying k-mer spectrum for a sample. We utilise consistent weighted sampling to incorporate k-mer frequency information into the histosketch, allowing the use of weighted and standard Jaccard similarity for histosketch comparisons and sample retrieval (Ioffe, 2010). Our method combines the recently proposed histogram sketching algorithm of Yang *et al.* with count-min sketching of k-mer spectra and our recent implementation of LSH Forest indexing for microbiome searching (Yang *et al*., 2017; Zhang *et al*., 2014; Rowe and Winn, 2018). We show our method to accurately cluster microbiome samples by sample type and demonstrate the utility of these histosketches to create and search microbiome sequence databases. Finally, we show that histosketches are suitable features for training ML classifiers and can accurately classify microbiome samples according to antibiotic treatment history in at-risk preterm infant populations.

## 2. Materials and Methods

Here we describe our method for the compact representation of microbiome sequencing data using similarity-preserving histosketches of streaming k-mer spectra (Figure 1.). We then document our implementation, HULK, and describe several use cases.

**Figure 1.**
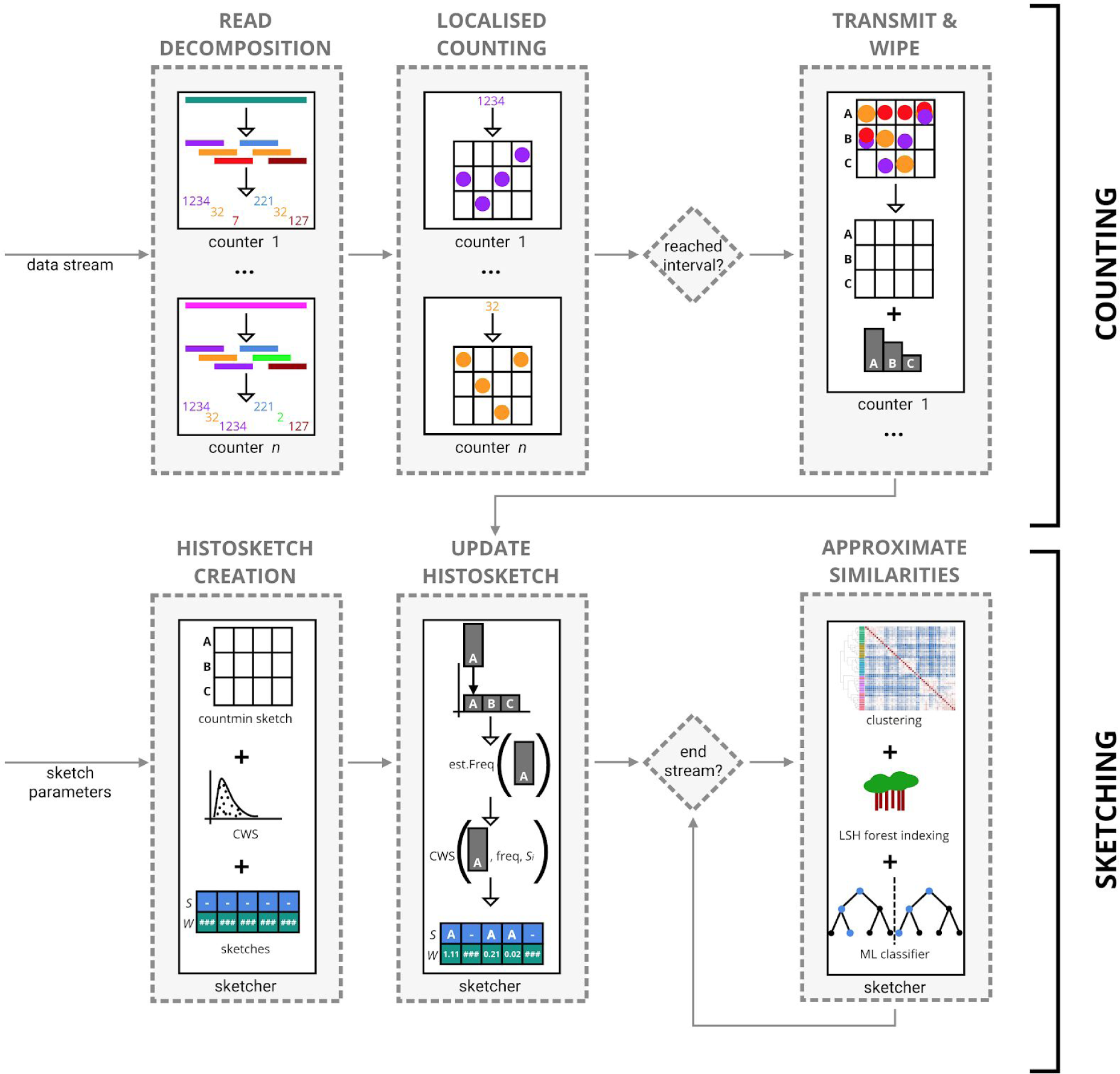
Overview of our method to histosketch microbiome samples from sequence data streams. **A.** During counting, sequence reads are collected from the data stream by *n* counting processes. Reads are decomposed to canonical k-mers, encoded to uint64 values and used to increment local countmin sketches. Once *X* reads have been received from the data stream, approximate k-mer counts from the counting processes are transmitted as histogram elements to the single sketching process. **B.** To update the histosketch, the incoming histogram element is hashed and compared against each hash value (*W*) or the previous histosketch (*S*), updating *S* and *W* if a new minimum is encountered. To hash the incoming vector, uniform scaling is applied and a cumulative frequency estimate is made using a countmin sketch; we then utilise CWS to generate a hash value for the updated histogram bin.

### 2.1. Histosketching microbiome data

We use the k-mer spectrum (a normalised vector of k-mer frequencies) to represent microbiome diversity, which is a standard analysis method that allows for metagenome dissimilarity analysis (Dubinkina *et al*., 2016; Benoit *et al*., 2016). However, rather than computing and storing a full k-mer spectrum after reading the sequence data, which is resource intensive (in terms of memory or disk space), we use the recently proposed histosketch data structure to maintain a set of fixed size sketches to approximate the overall k-mer spectrum as it is received from a data stream (Yang *et al*., 2017). The histosketch has two properties making it suitable for this application, *i*. it is updateable, and *ii*. it is similarity-preserving. Thus, as new data is received, we can incrementally update the histosketch of the underlying k-mer spectrum and also approximate similarity to other spectra.

We view the k-mer spectrum as a histogram, where k-mers from a microbiome sample are hashed uniformly across N bins and the frequency value of a bin corresponds to observed k-mer frequency. In order to incorporate both the bin and frequency (a weighted set) into the histosketch, we employ Consistent Weighted Sampling (CWS) to generate hash values for each histogram element, which ensures that the computational complexity of hashing is independent of bin frequency (Ioffe, 2010; Yang *et al*., 2017).

#### 2.1.1. Consistent Weighted Sampling

As highlighted in the introduction, a drawback to the efficient set similarity estimations afforded by MinHash sketches is that the input is restricted to binary sets and does not account for weighted sets (e.g. k-mer frequencies). To overcome this, histosketching employs CWS to account for element frequency and approximate the generalised Jaccard similarity between weighted sets, without splitting each weighted element into sub-elements and computing independent hash values (quantization) (Haveliwala *et al*., 2000; Manasse *et al*., 2010; Ioffe, 2010; Wu *et al*., 2017).

For a weighted set of k-mers, W, where k-mer frequency W_k_ ≥ 0 for all elements of the set, CWS produces a sample, (k, y_k_) : 0 ≤ y_k_ ≤ W_k_, which is both uniform and consistent. The sample is uniformly sampled from ∪_k_{k} × [0, W_k_], meaning that the probability of selecting k is proportional to the k-mer frequency, W_k_, and y is uniformly distributed on [0, W_k_]. The sample is also consistent as given two weighted sets, W1 and W2, if ∀k, W1_k_ ≤ W2_k_, a subelement (k, y_k_) is selected from W1 and satisfies y_k_ ≤ W2_k_, then (k, y_k_) will also be selected from W2 (Ioffe, 2010; Wu *et al*., 2017).

In order to generate a weighted MinHash code (y_k_, y_a_) for a member of a weighted set (W_k_), CWS uses two equations (Eq. 1 and Eq. 2), where r_k_ ∼ Gamma(2, 1), β_k_ ∼ Uniform(0, 1) and c_k_ ∼ Gamma(2, 1):

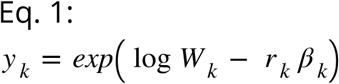

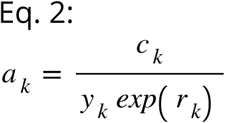

Eq. 1 generates an “active index” that is then fed in to Eq. 2 so that k is sampled in proportion to its sample weight. Eq. 2 outputs a hash value the conforms to the exponential distribution that is parameterised with the sample weight of W_*k*_.

#### 2.1.2. Histosketch creation

Equations 1 and 2 describe the CWS method, which we apply to sample a k-mer spectrum in a way that takes the relative abundance of k-mers into account. To generate a sketch of a k-mer spectrum originating from a biological sample, the k-mer spectrum is sampled Z times, where Z is the size of the sketch. We will denote our underlying k-mer spectrum (a histogram) as V, with cardinality |ε| = X (i = 1,…, X). The corresponding histosketch we will denote as S, with cardinality |ε| = Z (j = 1,…, Z). To initialise S from V, first three independent variables are sampled from the CWS distributions: r_i_,_j_ ∼ Gamma(2, 1), c_i_,_j_ ∼ Gamma(2, 1) and β_i_,_j_ ∼ Uniform(0, 1) for i ∈ E and j = 1,…, Z. We then use Algorithm 1 of Yang et al. for histosketch creation (Yang *et al*., 2017). The sketch, S, and the corresponding hash values, A, are both kept as the histosketch (A allows for incremental sketch updating).

##### Algorithm 1: Histosketch creation

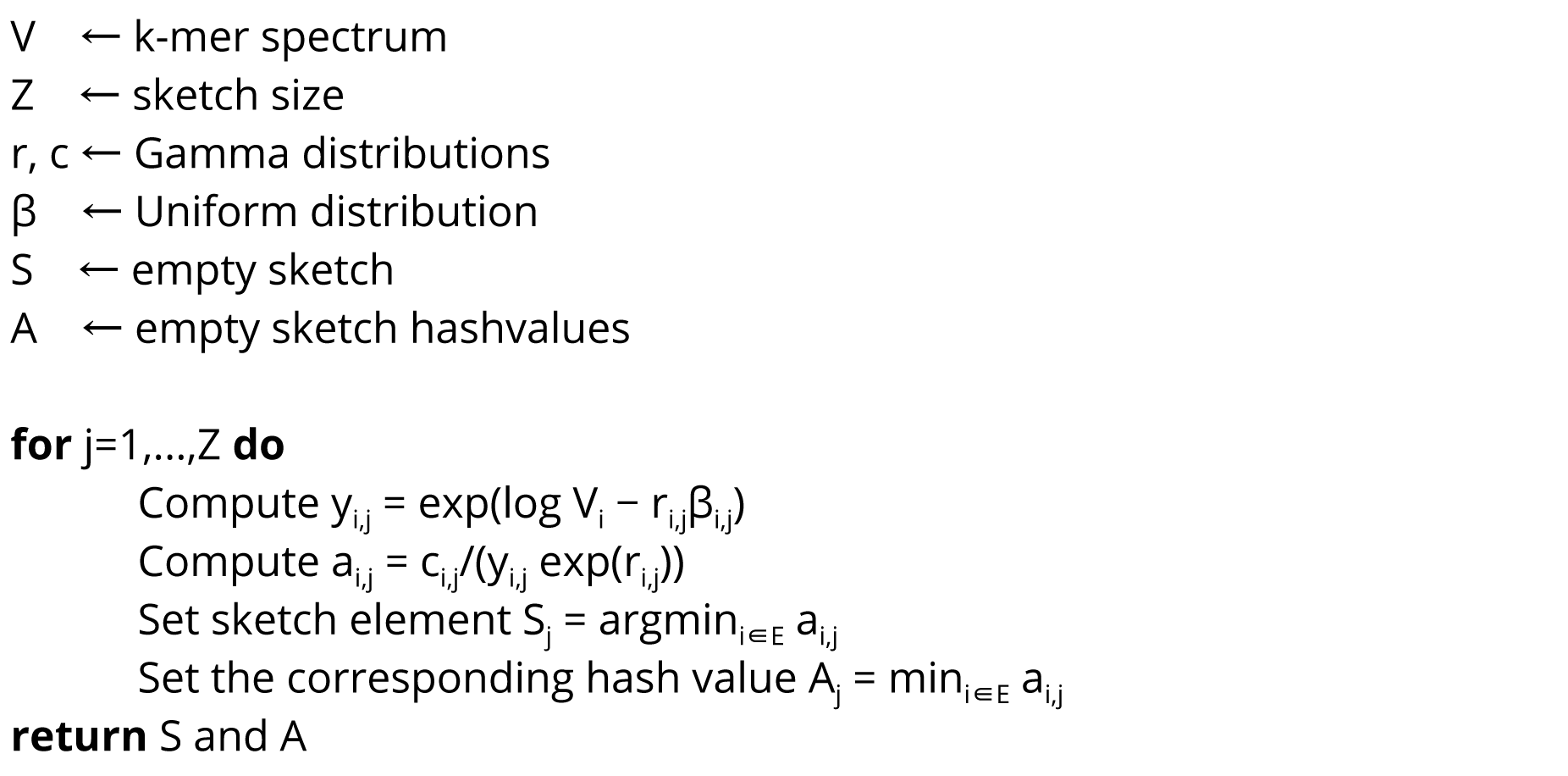

To summarise, to create an element (j = 1,…, Z) for one histosketch slot, based on underlying histogram V, we select the histogram element (i = 1,…, X) whose hash value is minimal and also keep the corresponding hash value.

#### 2.1.3. Histosketch updating

To update the histosketch as a new histogram element is received, the previous sketch and the sketch hash values (S and A) are required. In its simplest form, the histosketch incremental update works by hashing and evaluating the incoming element against each slot of the histosketch. The cumulative bin frequency of the incoming element is estimated using a persistent countmin sketch (Cormode and Muthukrishnan, 2005); the frequency estimate is then used to update the hash value for the required histogram bin. If this hash value is now a minimum, the sketch slot and corresponding hash value are updated.

In addition to this update method, we can also utilise the gradual forgetting weights of the original histosketch implementation to adjust for changes in the underlying distribution (concept drift) (Koychev, 2000; Yang *et al*., 2017). Prior to the update, uniform scaling is applied to the estimate frequency counts. After this, the histosketh hashes are scaled using a decay weight before evaluating against the incoming element.

### 2.2. Our implementation

We have implemented our method as an easy to use program called HULK. HULK is written in Go (version 1.11) and compiles for a variety of operating systems and architectures. It is also packaged for installation with Bioconda and Biocontainers (Björn Grüning *et al*., 2018; Bjorn Grüning *et al*., 2018). HULK utilizes a concurrent pipeline pattern that is driven by the flow of data between structs. This pattern facilitates the streaming of data from STDIN, as well as from disk, and allows the HULK subcommands to be piped together and operate on data streams.

#### 2.2.1. Histosketching

The HULK subcommand `sketch` performs histosketching on a FASTQ data stream. Reads are collected from the data stream by one or more independent counting processes (Figure 1: counting); each utilising a separate Go routine for concurrent counting. Each counting process will count reads until an interval is reached (e.g. 1 million reads have been seen) or a signal is sent (e.g. the sample has been classified using a downstream ML classifier, see section 2.2.4), the counting processes will then send their count data via a Go channel to be histosketched, then wipe their stores and collect more reads.

A read is received by the counting process as a slice of bytes and the canonical k-mers are encoded to unsigned integers (uint64) using bit shift operations. Once encoded, the k-mer frequency is updated in the local store of the counting process. To ensure the counting processes operate in a fixed amount of memory, we again use the countmin sketch data structure to record frequency estimates for the k-mer spectrum (Cormode and Muthukrishnan, 2005; Zhang *et* al., 2014). The countmin sketches are initialised with epsilon and delta values to control the relative accuracy and the resulting number of countmin counters are used as a proxy for the number of bins in the underlying kmer spectrum.

Once an interval is reached, the counting processes send their k-mer spectrum data in a randomised order to the single histosketching process; this process follows the incremental histosketch update process described above (Figure 1: sketching).

#### 2.2.2. Distance estimation

HULK includes two distance subcommands, `distance` and `smash`. Running `hulk distance` will run a pairwise comparison of two histosketches and output the Jaccard, weighted Jaccard, Bray Curtis or Euclidean metrics. Running `hulk smash` will perform a pairwise comparison of two or more histosketches and output a matrix of Jaccard or weighted Jaccard similarities. The calculation of weighted Jaccard distance utilises the histosketch bin and corresponding hash values; Eq. 3 shows the calculation of weighted Jaccard distance for two histosketches, S and T.

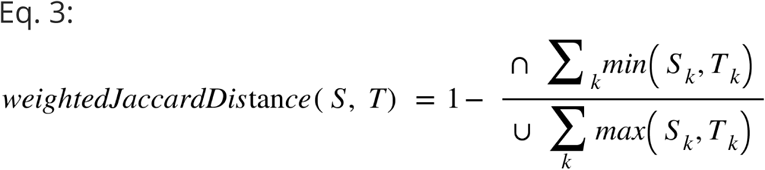

#### 2.2.3. Indexing

HULK utilises the LSH Forest self-tuning indexing scheme as employed in our previous work (Rowe and Winn, 2018). Briefly, this scheme will take a query and return a subset of nearest-neighbour candidates, based on the number of hash collisions (Bawa *et al*., 2005). The two parameters to tune this index are (i) the number of hash functions to encode an item (K) and (ii) the number of hash tables to split an item in to (L). To tune index prior to adding items, multiple combinations of K and L are evaluated by false positive/negative rate at the given Jaccard similarity threshold. To add a histosketch to the index we use only the sketch S (i.e. not the hash values A, see section 2.1.2); the sketch is split in to L equally sized chunks of K hashes. The chunks are hashed to a binary string (little-endian ordering) and stored in the corresponding hash table. Prior to searching the index, the hash tables are transferred to a set of arrays and sorted.

The HULK index operations are performed using the ‘index’ subcommand. Three modes are available: create, add and search. To create an index, the LSH Forest index is initialised using a Jaccard similarity and error rate thresholds, then each histosketch is split in to the appropriate number of chunks and added as described earlier. The index is written to disk in the unsorted form. To add a histosketch to an existing index, the index is loaded and the histosketch is added using the existing index parameters.

To search the index, the index is first loaded and the hash tables are transferred to a set of arrays and sorted. The query set of histosketches are then queried in series and the similar histosketches are returned (by label) that are within the Jaccard similarity threshold that was set during indexing.

#### 2.2.4. Random Forest Classifier

We implemented a Random Forest Classifier (RFC) as an example ML classifier to showcase the applicability of our histosketches as features for predicting microbiome sample labels. Our implementation (BANNER) is written in Python (version 3.6) and is distributed with HULK, as well as through Bioconda and Pypi. Source code is available at https://github.com/will-rowe/banner. It uses the SciKit Learn (version 0.19.2) implementation of the RFC (Pedregosa *et al*., 2011). Again, we use only the sketch values S and discard the hash values A. BANNER trains on 80% of the available data using bootstrapping and 1000 estimators; testing then uses the remaining 20% of the available data and does this with 10-fold cross validation. Once trained, the RFC model is serialised. To classify histosketches with BANNER, the RFC model is first loaded and un-serialised, before collecting histosketches from STDIN, allowing the output of `hulk sketch` to be piped so that histosketches can be classified as they are generated:

hulk sketch -f sample.fastq --stream -p 8 | banner predict -m banner.rfc

The predict subcommand will only terminate once it makes a prediction above a set probability threshold or the sketching processes finishes.

### 2.3. Evaluating performance

The full commands and code used to evaluate the performance of our implementation can be found in the HULK repository (https://github.com/will-rowe/hulk/tree/master/paper). HULK version 0.0.2 was used in all experiments (release 0.0.2, commit 97ba8ac).

For running the clustering and indexing experiments, the simulated short reads from the Critical Assessment of Metagenome Interpretation (CAMI) project (dataset to benchmark new programs against highly complex and realistic metagenomic datasets) were downloaded in FASTQ format (Sczyrba *et al*., 2017). For each read set, HULK sketches (k-mer size=21, histosketch size=512) and Simka (version 1.4.0) k-mer spectra (k-mer size=21) were created and pairwise Jaccard distances were loaded into Python (version 3.6.5) using Pandas (version 0.23.4) (Mc Kinney) and clustered using Seaborn (version 0.9.0) (clustering method=complete). For running HULK and Simka, both were restricted to 12 CPUs per FASTQ file and run using LSF on a high performance computing cluster (Atos Bull Sequana, Intel Skylake nodes).

As an additional clustering experiment, we used a recently published dog microbiome dataset to detect dietary intervention using histosketches on varying levels of sequencing data (ENA: PRJEB20308) (Coelho *et al*., 2018). This study reported a significant shift in the taxonomic composition of dog microbiomes when diets were changed from a baseline diet. The full dataset contains 1.9 terabasepairs of sequencing data, of which we sampled 0.005%, 0.05% and 0.5% of each microbiome. We histosketched these samples (k-mer size=21, histosketch size=512) and clustered them as above.

For performing the RFC analysis, an RFC model was constructed as described in 2.2.4, using a clinically relevant dataset; gut microbiome profiles from a cohort of healthy pre-term neonates from a single hospital. This is part of a wider clinical study, that is longitudinally profiling the gut microbiota of preterm infants that are residing in neonatal intensive care units (NICUs) and correlating this to health data, including impact of antibiotics. Faecal samples from preterm infants were collected and their bacterial DNA extracted following the protocols described in Alcon-Giner et al., 2017 (Alcon-Giner *et al*., 2017). Shotgun metagenomics libraries were prepared from 500 ng of genomic DNA which was sheared into fragments of ∼450 bp. The sheared DNA was purified and concentrated using an SPRI-clean-up kit. Library construction entailed an end repair, A-tailing and adapter ligation steps. Following, adapter ligation, samples were amplified and indexed by PCR using established Illumina paired end protocols. A portion of each library was used to create an equimolar pool and pooled libraries were subjected to 125 bp paired end sequencing on a HiSeq 2500 V4. The cohort was labelled according to whether the infants were receiving prophylactic antibiotic treatment or no antibiotics. The histosketches from 108 FASTQ files (BioProject: PRJEB28428) were split into training (80%) and testing (20%) groups. When using the RFC model to classify the incremental sketch updates of blinded samples, HULK was run using a sketching interval of 1,000,000 reads using a 4 core laptop (k-mer size=21, histosketch size=512).

## 3. Results

The results presented here evaluate our implementation of histosketching for rapid microbiome comparisons, in terms of both the accuracy of the tool and its potential applications. All analyses can be run using the analyses workbooks (https://github.com/will-rowe/hulk/tree/master/paper/analysis-notebooks)

### 3.1. Clustering microbiome datasets

The CAMI metagenome sequence data for 48 microbiome samples were sketched by HULK in 1 minute 30 seconds and the full k-mer spectra were computed by Simka in 24 minutes and 1 seconds. Hierarchical clustering identified 5 distinct groups using both the HULK histosketches (Figure 2a) and the full k-mer spectrum (Figure 2b). These groups corresponded to the 5 body sites of the CAMI project, as denoted by the coloured bars on the dendrograms. Using the HULK sketches, 2 samples failed to cluster by body site (skin and airways), whereas 3 samples failed to cluster for the full k-mer spectra (skin and airways).

**Figure 2.**
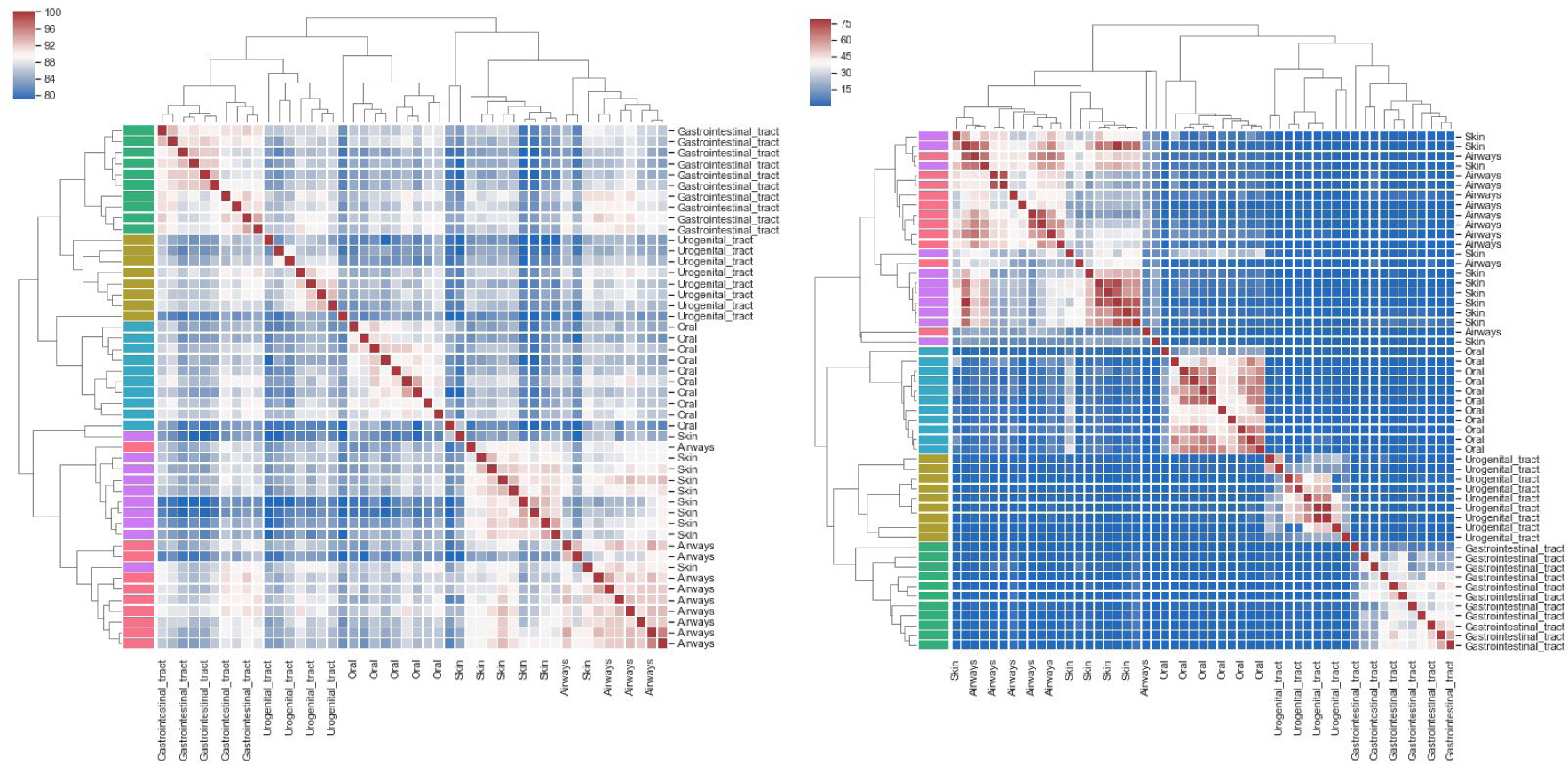
Hierarchical clustering of CAMI short read microbiome samples (Sczyrba *et al*., 2017). Heatmaps show the pairwise Jaccard similarity between microbiome samples; colormap ranges are computed using robust quantiles and dendrogram clades are coloured by body site. **A**. HULK histosketches (k-mer size=21, histosketch size=512) for 48 microbiome samples were sketched in 1 minute 30 seconds (12 cores per histosketch). **B**. Simka k-mer spectra (k-mer size=21) for 48 microbiome samples were computed in 24 minutes 1 seconds (12 cores per spectrum).

To show the ability of our method to cluster incomplete data streams in a biological meaningful way, we performed incremental histosketch updating on data streams from a collection of dog microbiome samples. As the data was downloading, we histosketched the data stream (using fastq-dump to stream the download); approximately 0.005%, 0.05% and 0.5% of the reads from each sample (129 samples total) were processed and then clustered based on pairwise Jaccard similarity (Figure 3). At all intervals, we found clear separation of histosketches between microbiome samples from dogs receiving the baseline diet and those receiving an altered diet (high/low protein). This is in agreement with the findings of the original study, where they reported a significant shift in the taxonomic composition of dog microbiomes when diets were changed (Coelho *et al*., 2018). The total microbiome data for the original study was stored in 3096 runs across 129 samples, amounting to 1.9 terabasepairs. Complete download of this dataset from the ENA took over 7 days using fastq-dump with 20 parallel downloads. Sketching the initial 0.005% of the data stream took an average of 4 seconds per sequencing run (approximately 100 seconds per sample).

**Figure 3.**
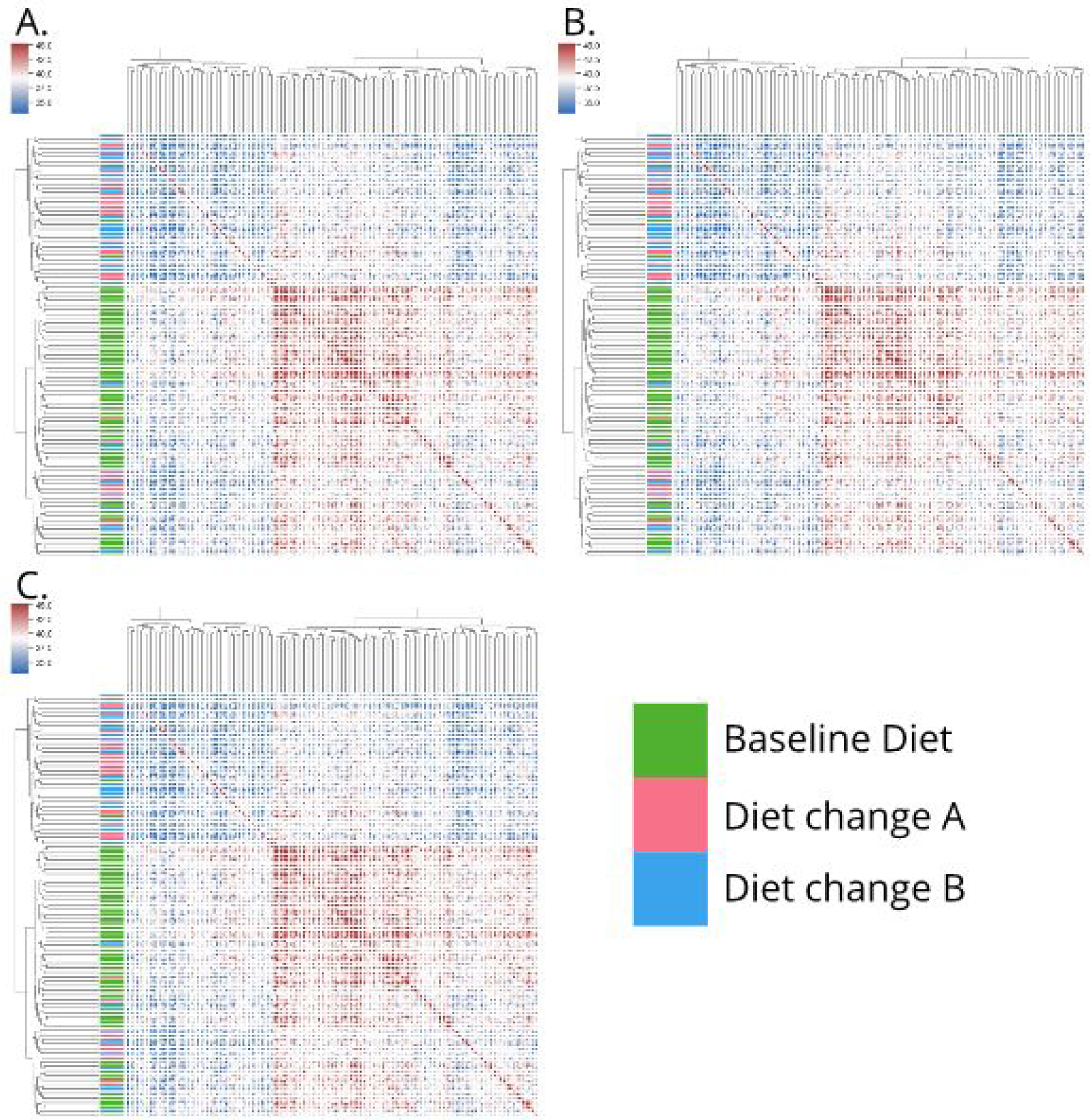
Hierarchical clustering of dog microbiome samples (Coelho *et al*., 2018). A, B and C correspond to clustered histosketches from 0.005%, 0.05% and 0.5% of sample reads respectively. The majority of microbiome samples from the dogs on the baseline diet clustered together (green), however the samples taken after these dogs were put on to an altered diet (pink/blue) did not show any distinct clustering pattern.

### 3.2. Indexing microbiome collections

The histosketches from the CAMI metagenome sequence data were labelled by body site before 1 sample was randomly removed from each group and used as search queries. The remaining sketches were indexed using HULK in 0.039 seconds (with a Jaccard similarity threshold of 0.90). All the queries returned at least one CAMI sample from the same body site (Figure 4). The oral query returned only oral samples, the Gastrointestinal (GI) tract, airways and skin queries returned predominantly samples from their own respective body sites, whilst the Urogenital (UG) tract returned one sample from the same body site, plus another from airways. When overlaid on Principal Components 1 and 2 of a PCA analysis, the search queries are grouped nearest their respective LSH Forest search results (Figure 4).

**Figure 4.**
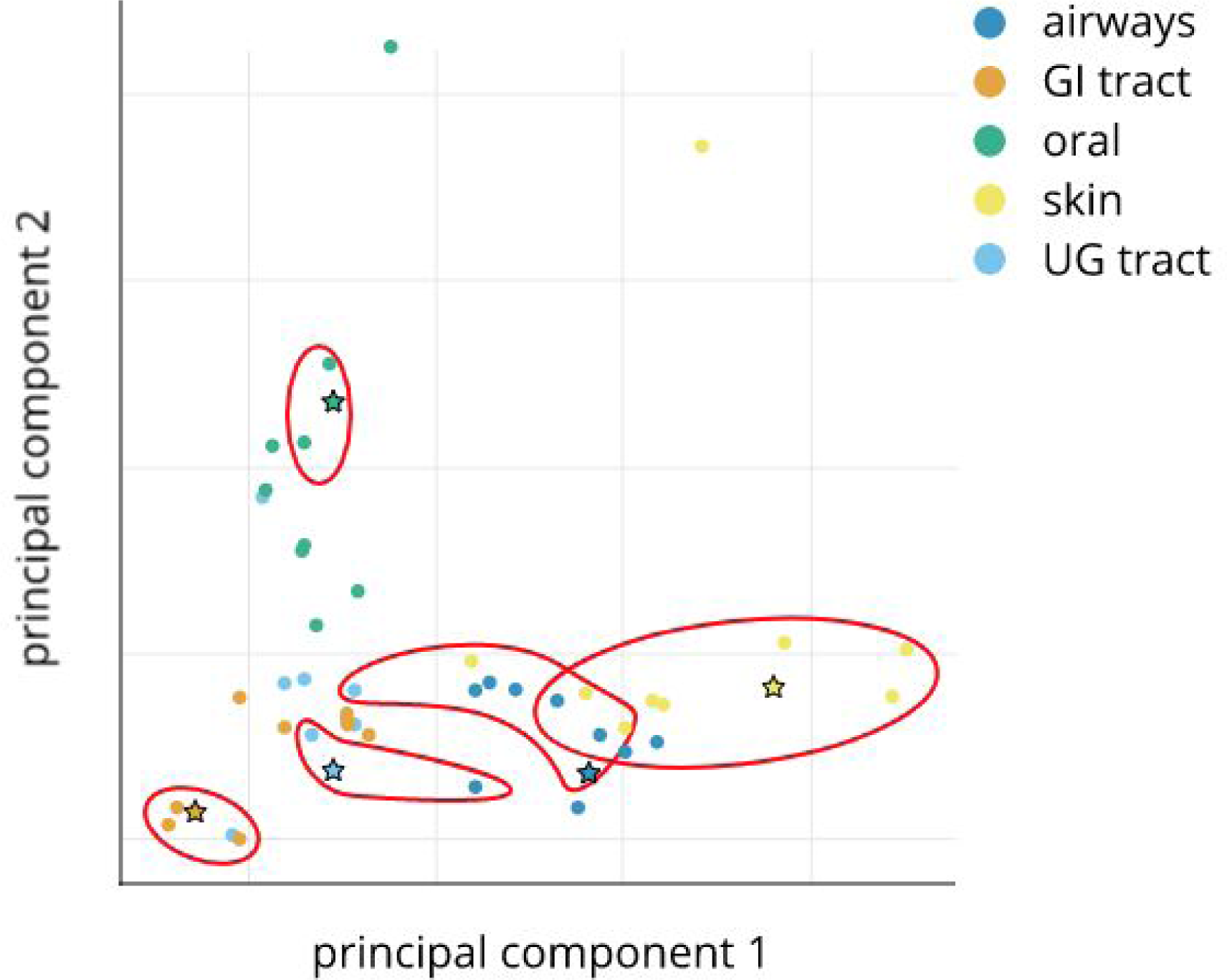
Principal component analysis of histosketches from CAMI short read microbiome samples, coloured by body site (Sczyrba *et al*., 2017). Circular data points indicate the histosketches used to build the LSH Forest index, stars data points indicate histoketches used as search queries. Red rings enclose the returned LSH Forest search results for each search query (Jaccard Similarity threshold > 90%).

### 3.3. Classifying microbiomes using machine learning

We trained a Random Forest Classifier using a clinically relevant microbiome collection (gut microbiome profiles from a cohort of healthy pre-term neonates from a single NICU) and labelled the samples according to whether the infants were receiving prophylactic antibiotic treatment or no antibiotics. The accuracy on the test set during RFC construction was 0.95, with an F1 score of 0.92 (Figure 5). When streaming reads from blinded microbiome samples from the cohort, histosketches after the first sampling interval (1,000,000 reads, average sketching time = 12 seconds) were then successfully classified by the previously trained RFC as being from an antibiotic treated neonate (probability = 0.90), upon which the data stream was terminated and a new sample data stream was then histosketched.

**Figure 5.**
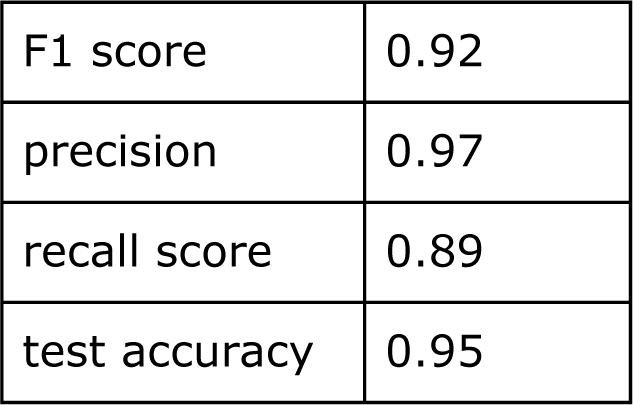
Random Forest Classification test accuracy for predicting antibiotic treated vs. no-antibiotic treated neonatal microbiomes.

## 4. Discussion

In this paper we have presented a new method, as well as several use cases, for rapid microbiome analytics using streaming histogram sketching. This work has been in direct response to the call for improved microbiome analytics in this era of big data, massive microbiome sequencing initiatives and the realistic prospect of clinical metagenomics (Kakkanatt *et al*., 2018; Mulcahy-O’Grady and Workentine, 2016). We feel that our microbiome sketching method and the applications shown here go toward addressing this challenge.

As outlined in the introduction, the dimensionality reduction methods that have only recently been applied to genomics have been a great advance toward the goal of rapid microbiome analytics; facilitating fast similarity queries such as identifying genomes or genes within metagenome samples (Brown and Irber, 2016; Rowe and Winn, 2018). Our dimensionality reduction method for the comparison, indexing and classification of microbiomes offers a novel and complementary method to these existing ones. In particular, it addresses the main limitations of traditional MinHash for certain microbiome analyses. These being: (i) histogram sketching is not impacted by mismatched set size (Koslicki and Zabeti, 2017) and (ii), histogram sketching accounts for weighted sets (e.g. k-mer frequency).

In terms of the advantages of our method over other *de novo* analysis methods (e.g. k-mer spectra dissimilarity analysis), we have shown here that the computation of histogram sketches is 16 times faster than computation of the full k-mer spectra (see Results 3.1). As well as faster analysis times, histogram sketching has a much smaller footprint as the entire k-mer spectrum does not need to be kept in memory or written to disk, and the resulting sketches are much smaller in size than the full spectrum. Our method also does not require re-computation of previously sketched samples in order to make new comparisons. In addition, we showed that histosketching microbiome samples can work on incomplete data streams and allow samples to be clustered by the underlying microbiome composition when using just a small proportion of the total reads (see Results 3.1, Figure 3). Our results suggest that only a small proportion of the total data stream needs to be sampled in order to cluster the samples according to a particular treatment using histosketch similarity. Although we managed to identify the time point when the diet was changed from baseline to an altered diet, we were not able to differentiate between the two altered diets using our sketches from the initial data stream. This may be due to insufficient sampling of the data stream; however, the original study did not report being able to differentiate between the two altered diets either (see figure 2c, Coelho et al. (Coelho *et al*., 2018)).

In sections 3.2 and 3.3, we demonstrated that microbiome histosketches can be efficiently indexed and also used as features in ML classification, which are both typically hard to do using the full k-mer spectra due to their scale and sparsity (Kakkanatt *et al*., 2018). In terms of the LSH Forest index for microbiome sample retrieval, our results showed that a histosketch from a given body site would predominantly return microbiome samples from the same body site (Figure 4). Only the oral histosketch query returned solely oral samples, which is likely due to the high similarity observed between these datasets (Figure 2). On the whole, these results indicate that histosketches of k-mer spectra can offer an efficient and fast way to index and query collections of microbiome data.

Our performance evaluation of HULK using an RFC illustrates how incremental sketching (as highlighted in Figure 3) can be combined with ML in order to classify a microbiome and stop processing a data stream (see Results 3.3). This is a step forward in dealing with streaming genomics data; the combination of incremental histosketch updates with a ML classifier (and associated classification probabilities) allows for the possibility of terminating data streams in applications such as real-time sequencing (Ondov *et al*., 2016). Here, we used this approach to quickly evaluate longitudinal samples from a cohort, identifying whether there is a response to a treatment or other stimulus. In this case we have used this sketching approach to differentiate between those preterm infants that had received antibiotics, versus those that did not. This is important clinically as antibiotic treatment in preterm infants is associated with significant alterations in the gut microbiota, which may link to increase risk of development serious conditions such as necrotising enterocolitis or sepsis (Sim *et al*., 2015; Shaw *et al*., 2015; Alcon-Giner *et al*., 2017). Thus a rapid and discriminatory microbiome profiling method for this fragile and at-risk patient cohort, or indeed for other clinical microbiome samples, could prove useful for intervention or treatment options. Alternatively, it could be applied to real-time sequencing platforms and inform the sequencer when enough data has been produced. These examples illustrate how this method could be used in the coming era of clinical metagenomics (Mulcahy-O’Grady and Workentine, 2016). We are not restricted to using RFC as and it would be very useful to evaluate other more sophisticated ML approaches that can utilise histosketches as feature vectors. As well as this, we could refine our ML models further by identifying the more significant elements of the histosketch in terms of their influence over the model training. This in turn could reveal more information relating to the underlying k-mer spectrum of a sample, which may be of use in downstream applications (e.g. feature extraction).

For future work into microbiome analytics, the histogram sketching method presented has potential for further refinement and improvements in order meet the big data challenges that microbiome research presents. Of these, we have already identified that further work into the use of histosketches in ML is definitely needed, particularly with the hope of improving classification accuracy and expanding out from the binary classification task we have shown here. In addition, we would like to explore the idea of concept drift for gradually forgetting outdated histogram elements (Yang *et al*., 2017). We included the gradual forgetting of the original histosketch method in our implementation but this was not exploited in the performance analysis (Yang *et al*., 2017). We envisage that this could be useful to experiment with in terms of real-time sequencing applications. Finally, we have shown that microbiome samples can be histosketched on a laptop with a few cores and a small, fixed amount of memory. In order to fully take advantage of this performance, histosketching needs to move beyond command line interfaces. To this end, we have begun work on a WebAssembly (WASM) port of HULK to enable client side sketching (WASM available Go Version 1.11) so that users can histosketch their own microbiome data and compare just the sketches against online databases, ensuring their microbiome data remains private but enabling quick and easy microbiome analytics.

## 5. Conclusions

To conclude, histosketching generates compact representations of microbiomes from data streams; facilitating sample indexing, similarity-search queries, clustering, and the application of machine learning methods to analyse microbiome samples in the context of the global microbiome corpus.

## Availability and implementation

The source code for our implementation, as well as the code used to run the analyses and plot the manuscript figures, can be found in the HULK (github.com/will-rowe/hulk) and BANNER (github.com/will-rowe/banner) repositories (MIT Licenses, DOIs: <u>10.5281/zenodo.1406952</u>, <u>10.5281/zenodo.1406951</u>).

## Authors’ contributions

W.P.M.R. and A.P.C conceived and executed the study. W.P.M.R coded and tested the implementation. LJH and JSK led on the preterm clinical study, and AS, KS, and CAG processed and sequenced samples, and SC carried out metagenomics QC, processing and analysis. All authors wrote, read and approved the final manuscript.

## References

Alcon-Giner, C. et al. (2017) Optimisation of 16S rRNA gut microbiota profiling of extremely low birth weight infants. BMC Genomics, 18, 841.

Anvar, S.Y. et al. (2014) Determining the quality and complexity of next-generation sequencing data without a reference genome. Genome Biol., 15, 555.

Bawa, M. et al. (2005) LSH Forest: Self-tuning Indexes for Similarity Search. In, Proceedings of the 14th International Conference on World Wide Web, WWW ′05. ACM, New York, NY, USA, pp. 651–660.

Benoit, G. et al. (2016) Multiple comparative metagenomics using multiset k-mer counting. PeerJ Comput. Sci., 2, e94.

Brown, T. and Irber, L. (2016) sourmash: a library for MinHash sketching of DNA. JOSS, 1, 27.

Coelho, L.P. et al. (2018) Similarity of the dog and human gut microbiomes in gene content and response to diet. Microbiome, 6, 72.

Cormode, G. and Muthukrishnan, S. (2005) An improved data stream summary: the count-min sketch and its applications. J. Algorithm. Comput. Technol., 55, 58–75.

Dubinkina, V.B. et al. (2016) Assessment of k-mer spectrum applicability for metagenomic dissimilarity analysis. BMC Bioinformatics, 17, 38.

Forbes, J.D. et al. (2018) Highlighting Clinical Metagenomics for Enhanced Diagnostic Decision-making: A Step Towards Wider Implementation. Comput. Struct. Biotechnol. J., 16, 108–120.

Greninger, A.L. et al. (2015) Rapid metagenomic identification of viral pathogens in clinical samples by real-time nanopore sequencing analysis. Genome Med., 7, 99.

Grüning, B. et al. (2018) Bioconda: sustainable and comprehensive software distribution for the life sciences. Nat. Methods, 15, 475–476.

Grüning, B. et al. (2018) Recommendations for the packaging and containerizing of bioinformatics software. F1000Res., 7.

Haveliwala, T. et al. (2000) Scalable Techniques for Clustering the Web (Extended Abstract).

Human Microbiome Project Consortium (2012) Structure, function and diversity of the healthy human microbiome. Nature, 486, 207–214.

Ioffe, S. (2010) Improved Consistent Sampling, Weighted Minhash and L1 Sketching. In, 2010 IEEE International Conference on Data Mining., pp. 246–255.

Kakkanatt, C. et al. (2018) Curating and integrating user-generated health data from multiple sources to support healthcare analytics. IBM J. Res. Dev., 62, 2:1–2:7.

Koslicki, D. and Zabeti, H. (2017) Improving Min Hash via the Containment Index with applications to Metagenomic Analysis. bioRxiv, 184150.

Koychev, I. (2000) Gradual Forgetting for Adaptation to Concept Drift. In, In Proceedings of ECAI 2000 Workshop Current Issues in Spatio-Temporal Reasoning.

Libbrecht, M.W. and Noble, W.S. (2015) Machine learning applications in genetics and genomics. Nat. Rev. Genet., 16, 321–332.

Luo, Y. et al. (2018) Metagenomic binning through low-density hashing. Bioinformatics.

Manasse, M. et al. (2010) Consistent Weighted Sampling. Financial Times.

Mc Kinney, W. pandas: a Foundational Python Library for Data Analysis and Statistics.

Morgan, X.C. and Huttenhower, C. (2012) Chapter 12: Human microbiome analysis. PLoS Comput. Biol., 8, e1002808.

Mulcahy-O’Grady, H. and Workentine, M.L. (2016) The Challenge and Potential of Metagenomics in the Clinic. Front. Immunol., 7, 29.

Ondov, B.D. et al. (2016) Mash: fast genome and metagenome distance estimation using MinHash. Genome Biol., 17, 132.

Pedregosa, F. et al. (2011) Scikit-learn: Machine Learning in Python. J. Mach. Learn. Res., 12, 2825–2830.

Rowe, W.P.M. and Winn, M.D. (2018) Indexed variation graphs for efficient and accurate resistome profiling. Bioinformatics.

Rusch, D.B. et al. (2007) The Sorcerer II Global Ocean Sampling expedition: northwest Atlantic through eastern tropical Pacific. PLoS Biol., 5, e77.

Sczyrba, A. et al. (2017) Critical Assessment of Metagenome Interpretation-a benchmark of metagenomics software. Nat. Methods, 14, 1063–1071.

Seth, S. et al. (2014) Exploration and retrieval of whole-metagenome sequencing samples. Bioinformatics, 30, 2471–2479.

Shaw, A.G. et al. (2015) Late-Onset Bloodstream Infection and Perturbed Maturation of the Gastrointestinal Microbiota in Premature Infants. PLoS One, 10, e0132923.

Sim, K. et al. (2015) Dysbiosis anticipating necrotizing enterocolitis in very premature infants. Clin. Infect. Dis., 60, 389–397.

Thompson, L.R. et al. (2017) A communal catalogue reveals Earth’s multiscale microbial diversity. Nature, 551, 457–463.

Wu, W. et al. (2017) Consistent Weighted Sampling Made More Practical. In, Proceedings of the 26th International Conference on World Wide Web. International World Wide Web Conferences Steering Committee, pp. 1035–1043.

Yang, D. et al. (2017) HistoSketch: Fast Similarity-Preserving Sketching of Streaming Histograms with Concept Drift. In, 2017 IEEE International Conference on Data Mining (ICDM)., pp. 545–554.

Zhang, Q. et al. (2014) These are not the k-mers you are looking for: efficient online k-mer counting using a probabilistic data structure. PLoS One, 9, e101271.

